# Impact of social isolation stress on safety learning and the structure of defensive behavior during a spatial-based learning task involving thermal threat

**DOI:** 10.1101/2024.09.28.615577

**Authors:** Stephanie A. Villalon, Ada C. Felix-Ortiz, Kelly Lozano-Ortiz, John R. McCarrey, Anthony Burgos-Robles

## Abstract

Safety learning during threat and adversity is critical for behavioral adaptation, resiliency, and survival. Using a novel mouse paradigm involving thermal threat, we recently demonstrated that safety learning is highly susceptible to stress. Yet, our previous study primarily considered male mice and did not thoroughly scrutinize the relative impacts of stress on potentially distinct defensive mechanisms implemented by males and females during the thermal task. The present study assessed these issues while analyzing a variety of defensive behaviors related to safety-seeking, escape, coping, protection, ambivalence, and risk-taking. After a two-week social isolation stress period, mice were required to explore a box arena that had thermal threat and safety zones (5°C vs 30°C, respectively). Since visuospatial cues clearly differentiated the threat and safety zones, the majority of the no-stress controls (69-75%) in both sexes exhibited optimal memory formation for the safety zone. In contrast, the majority of the stress-exposed mice in both sexes (69-75%) exhibited robust impairment in memory formation for the safety zone. Furthermore, while the control groups exhibited many robust correlations among various defensive behaviors, the stress-exposed mice in both sexes exhibited disorganized behaviors. Thus, stress severely impaired the proper establishment of safety memory and the structure of defensive behavior, effects that primarily occurred in a sex-independent manner.

## Introduction

Safety learning shapes behavior during threat and adversity to prevent harm and promote resiliency. While this function enhances adaptation and survival, deficiencies in safety learning in humans have been associated with profound alterations in behavior in stress-related disorders (Jovanovic et al., 2012; Laing and Harrison, 2021; Friedman et al., 2011). However, the mechanisms by which stress affects safety learning still remain elusive and highly neglected, while more attention has been given to how stress affects other related phenomena, such as facilitation of threat learning, impairment of cue and context discrimination, and failure in extinction learning (Conrad et al., 1999; Baratta et al., 2007; Holmes and Wellman, 2009; Dunsmoor and Paz, 2015; Maren and Holmes, 2016; Asok et al., 2018).

In a recent study, we implemented a novel approach to demonstrate that safety learning is highly susceptible to stress (Felix-Ortiz et al., 2024). Briefly, after undergoing social isolation stress for nearly two weeks, mice were exposed to a box arena containing multiple noxious cold quadrants (5°C, “threat zones”) and a more comfortable warm quadrant (30°C, “safety zone”). While visual cues differentiated these quadrants, ∼83% of the mice in the control group exhibited optimal memory for the thermal safety zone, whereas ∼67% of the mice in the stress group exhibited robust impairment of memory for the safety zone. Despite this striking finding, other important issues were not examined in our previous study. First, we did not sufficiently scrutinize the impacts of stress on stereotypical defensive behavior (Blanchard et al., 1986; LeDoux and Daw, 2018). Second, the previous study primarily considered male mice. Therefore, further investigation is needed to dissect potentially distinct impacts of stress in males and females on safety-seeking and defensive behavior (Luine et al., 2017; Merz and Wolf, 2017).

Emerging evidence suggests that male and female rodents could implement distinct behavioral strategies to deal with threat and adversity (Shansky, 2018, 2024; Foilb et al., 2021; Krueger and Sangha, 2021). Furthermore, female rodents often show more susceptibility to stress than males, especially during paradigms involving threat (Velasco et al., 2019; Day and Stevenson, 2020; Sangha et al., 2020; Furman et al., 2022). For instance, females exhibit stronger contextual threat conditioning and context generalization than males (Keiser et al., 2017). Females also exhibit more cue generalization than males during threat conditioning tasks involving CS+/CS-discrimination (Day et al., 2016). These observations are consistent with the fact that human female subjects exhibit greater risk for developing stress and trauma-related disorders (Kessler et al., 2012; Bangasser and Valentino, 2014; Shansky, 2015).

The present study implemented a comparative design with balanced samples for male and female mice to examine stress-related alterations in safety-seeking and stereotyped defensive behavior. While mice were subjected to our spatial-based thermal task after social isolation, three major categories were considered for defensive behavior. The first category included active defensive mechanisms such as rearing, darting, and jumping, which represent escape-related behaviors (Bolles, 1969; Gruene et al., 2015). The second category included passive defensive mechanisms such as freezing, crouching, and stretched posture, which represent protection, coping, and ambivalence, respectively (Van der Poel, 1967; Blanchard and Blanchard, 1969). The third category included other behaviors of relevance, such as grooming, general locomotion, and risk-taking (Spruijt et al., 1992; Ferreira et al., 2022). In summary, while stress altered safety memory and the relationships of safety-seeking behavior with other defensive mechanisms, there were no major sex differences in these effects.

## Materials and Methods

### Subjects

A total of 64 C57BL6/J young adult mice (10-weeks-old, 32 males and 32 females) were obtained from a commercial source (The Jackson Laboratory). Upon arrival to our local vivarium, mice were housed in groups of four with food and water ad libitum, in a room with regulated temperature, pressure, and a regular 12-hr light-dark cycle. After a week of acclimation, mice were brought into the laboratory space to undergo ear punching for identification and multiple handling sessions to become accustom to human interactions. All animal procedures were conducted after approval by the Institutional Animal Care and Use Committee, in accordance with the Guide for the Care and Use of Laboratory Animals and the U.S. Public Health Service’s Policy on Humane Care and Use of Laboratory Animals.

### Social Isolation Stress

A social isolation procedure was implemented to evaluate the impact of stress on safety learning (Felix-Ortiz et al., 2024). Social isolation represents a major non-invasive form of stress that is capable of producing profound impacts on mood, affect, cognition, and behavior (Ieraci et al., 2016; Mumtaz et al., 2018; Lee et al., 2021). For this, individual cages containing four mice each (all males or all females) were randomly assigned to the control or stress groups. The no-stress controls (“CTL”) were always kept in group-housing conditions, whereas the social isolation stress groups (“SIS”) underwent single-housing conditions for a period of 14 days, after which they were regrouped with their original cagemates. Bodyweights were recorded every other day to evaluate the effectiveness of social isolation as a stressor that impairs weight gain (Arias et al., 2010; Mumtaz et al., 2018). Further validation of the social isolation stress procedure was achieved through assessment in the elevated-plus maze and open-field assays, which often reveal stress-induced anxiety-like behaviors (Calhoon and Tye, 2015). The plus-maze and open-field assays were conducted one day before and one day after the social isolation procedure. Seven days after the end of social isolation, mice underwent training and testing in the thermal safety task.

### Elevated-Plus Maze

The elevated-plus maze test was used to evaluate the effectiveness of the social isolation stress treatment. This test was conducted in a plus-shaped apparatus that consisted of two open arms (L:30 cm × W:5 cm) and two enclosed arms (L:30 cm × W:5 cm × H:15 cm) that intersected at a small central zone (L:5 cm × W:5 cm). The apparatus was significantly elevated from the benchtop (H:40 cm) and constructed from a beige-colored ABS plastic material (San Diego Instruments). This assay was performed twice (10 min each time), one day before and one day after the social isolation treatment (i.e., pre-stress vs post-stress testing). Since this strategy could lead to reductions in general exploration due to decreased contextual novelty during the second test, the pre-stress and post-stress tests were conducted in distinct procedure rooms with distinct overall contextual settings (e.g., different odors, different light patterns, different lab coats worn by the student experimenters), in a counterbalanced fashion across mice to minimize such reductions. The time that mice spent in the open arms of the plus-maze apparatus was evaluated as a proxy of anxiety-like behavior (Walf and Frye, 2007). Significant reductions in open-arm exploration often emerge after stress exposure (Campos et al., 2013).

### Open-Field Test

The open-field test was performed to evaluate potential alterations in general locomotion by the social isolation stress treatment. The open-field assay was conducted in square-shaped transparent acrylic boxes (L:40 cm × W:40 cm × H:40 cm; Marketing Holders). Similar to the plus-maze assay, the open-field test was performed twice (10 min each time), one day prior to the beginning of social isolation and once again one day after the end of the social isolation treatment (i.e., pre-stress vs post-stress testing). To minimize reductions in exploratory behavior due to decreased contextual novelty, the pre-stress and post-stress tests were conducted in distinct procedure rooms with distinct odors and illumination patterns in a counterbalanced fashion across mice. The average speed of locomotion was quantified using software as mice navigated the open-field arena. Significant reductions in locomotor activity during the open-field test could represent alterations in exploratory behavior or depressed states, which are common after stress (Yan et al., 2010; Seibenhener and Wooten, 2015).

### Thermal Safety Task

The impacts of social isolation stress on safety learning and the structure of defensive behavior were evaluated using a spatial-based thermal threat task that we recently developed (Felix-Ortiz et al., 2024). The task consisted of exposing mice to a square-shaped arena in which three quadrants had a significantly noxious cold temperature (∼5°C, “threat zones”), whereas the remaining quadrant had a more comfortable warm temperature (∼30°C, “safety zone”). These temperatures are consistent with previous reports of adverse and preferred temperatures, respectively, in mice (Gordon et al., 1998; Bautista et al., 2007). The cold and warm temperatures were achieved by placing ice or handwarmers underneath the floor. The stability of the temperatures was monitored using a digital thermometer gun (LaserPro LP300; Kizen) and a thermal imaging camera (C5; FLIR). Readjustments to the temperatures were made in between animals as needed. The task was conducted in transparent acrylic boxes (L:30 cm × W:30 cm × H:30 cm; Marketing Holders). These boxes were placed inside sound-attenuating cabinets (L:70 cm × W:86 cm × H:56 cm; Med Associates), which were equipped with fans that provided constant airflow and background noise (65 dB) to reduce distractions from external noise. The cabinets also reduced the role of distal visual cues within the procedure room to modulate spatial learning, an effect shown in our previous study (Felix-Ortiz et al., 2024). The cabinets were also equipped with regular white lights, near-infrared lamps, and overhead infrared cameras for the recording of videos. Proximal visual cues were added to the walls of the acrylic boxes to facilitate spatial learning. For example, plus symbols were paired with the warm zone, whereas vertical bars were paired with the cold zones. These visuospatial cues were counterbalanced across mice. The task consisted of two sessions. The training session lasted 10 min and included the warm and cold zones. In contrast, the recall test (conducted 24 hr after training) lasted 5 min and only included a uniform cold temperature throughout the floor. The rationale for this was that if mice truly learned the precise location of the warm zone during training, the next day, they should still perform greater safety-seeking behavior within the correct zone even in the absence of the warm reinforcer, as validated in our previous study (Felix-Ortiz et al., 2024).

### Data Analysis

All behavioral sessions were recorded at 15 fps. Mouse behavior was later quantified using two methods. The first method consisted of automated tracking of mice using software (ANYmaze; Stoelting). This method was implemented to measure various behaviors during the thermal task, including (a) Safety-seeking behavior, which was defined as the total time that mice spent within the spatial zone that predicted the warm temperature; (b) Freezing behavior, defined as periods of minimal mobility for at least half a second; (c) Risk-taking behavior, defined as periods in which mice protruded the head into the threat zones while keeping the rest of the body within the safety zone; and (d) General locomotion, assessed as the average speed of motion exhibited during a given behavioral session. Software-based quantifications were also implemented during the elevated-plus maze to measure the amount of time that mice spent in the open arms of the apparatus and during open-field testing to measure locomotion.

The second method used for behavioral quantifications consisted of software-assisted continuous hand-scored sampling of behavioral events. This method is still regarded as a highly validated strategy to quantify animal behavior (Altmann, 1974; Oldfield, 2001; Hämäläinen et al., 2016; Brereton et al., 2022), and was implemented by student experimenters who were blind to mouse sex and treatment, similar to previous studies (Lehner, 1979; Felix-Ortiz et al., 2016). The following behaviors during the thermal task were quantified using this method: (a) Rearing behavior, defined as periods in which mice adopted an upright standing posture with their forelimbs touching the sidewalls of the box; (b) Darting behavior, defined as episodes in which mice suddenly made flight-like running moves; (c) Jumping behavior, which consisted of sudden hopping or leaping moves; (d) Crouching behavior, defined as periods in which mice adopted a hunched posture with the head and forelimbs rolled inwards while supporting the weight of the body on the hindlimbs; (e) Grooming behavior, defined as sequences of self-cleaning behavior; and (f) Body stretching, defined as periods of low mobility while mice exhibited an elongated body posture.

An important correction was made between two of the measurements. While crouching behavior was hand-scored, freezing behavior was scored by the software. These behaviors are somewhat related to each other and most likely were quantified together by the software (Blanchard and Blanchard, 1969; Anagnostaras et al., 2010). To distinguish these two measurements, the total number of crouching episodes was subtracted from the total number of freezing episodes to correct the latter quantification.

Raw data was extracted from the software (ANYmaze) and tabulated using spreadsheets (MS Excel). After data analysis, graphs were generated, and the results were further evaluated using statistical software (GraphPad Prism). Each behavior was compared across the male, female, no-stress, and stress groups, as well as across behavioral sessions. Due to differences in the length of the training and recall sessions (10 min for training and 5 min for recall), all behaviors were analyzed in normalized forms (e.g., percent time in the safety zone, event rates per minute, or average speed of motion). The normality of the data was verified using the Kolmogorov-Smirnov test. Consistently, all results were plotted as mean ± standard error of the mean, with values for individual mice illustrated as scattered plots over the bar graphs. Significant differences across groups and sessions were evaluated using two-way repeated measures analysis of variance (ANOVA), with Holm-Šídák post hoc tests for more statistical power (Salkind, 2007). Multiplicity-adjusted *P*-values were always considered to account for multiple comparisons. Chi-square tests were used to compare categorical proportions of mice that exhibited optimal memory versus poor memory, or stress-induced susceptibility versus stress resiliency (McHugh, 2013). Linear regression analysis was implemented to evaluate relationships among the distinct behaviors (Sheskin, 2003; Kleinhappel et al., 2019). For statistical significance, various two-tailed alpha levels were considered: \**P* < 0.05, \*\**P* < 0.01, \*\*\**P* < 0.001, or \*\*\*\**P* < 0.0001.

### Availability of Data and Materials

All the materials used in this study were obtained from commercial sources, as indicated throughout the methods. Software code and data will be shared for scientific use upon reasonable request.

## Results

### Overall experimental design to evaluate the effects of social isolation stress in safety learning

We recently showed that a two-week social isolation stress period was highly detrimental for the formation of lasting representations of safety when exposed to a box containing thermal threat and safety zones (Felix-Ortiz et al., 2024). Since our previous study primarily considered male mice, the first goal of the present study was to determine whether the effects of this stressor are sex-specific or sex-independent. We implemented a comparative design that included balanced samples of male and female mice (N = 32 for each sex). Initially, these mice were separated by sex and were housed in groups of four for approximately two weeks. Then, the cages were randomly assigned to either the control or experimental groups. While the control groups were kept in group-housing conditions (i.e., no-stress, CTL-M or CTL-F; N = 16 per sex), mice in the experimental groups underwent a two-week isolated-housing period (i.e., social isolation stress, SIS-M or SIS-F; N = 16 per sex). After the two-week isolation period, the stressed mice were regrouped with their former cagemates (Figure 1A). One week later, all mice underwent training in the thermal task in which the behavioral arena had three noxious cold zones and a comfortable warm zone (Figure 1B). The warm zone was differentiated from the cold zones using discrete visuospatial cues on the wall of the apparatus. As in our previous study, the most prominent behavior exhibited by mice during training was safety-seeking, which was evaluated as total time spent within the warm zone (Figure 1C). The next day, mice underwent a test session in the absence of the warm temperature to evaluate long-term recall of the safety memory (Figure 1D). During the recall test, based on the position of the visuospatial cues on the walls, biased seeking behavior towards the zone that previously predicted thermal safety was indicative of optimal memory (Felix-Ortiz et al., 2024).

**Figure 1.**
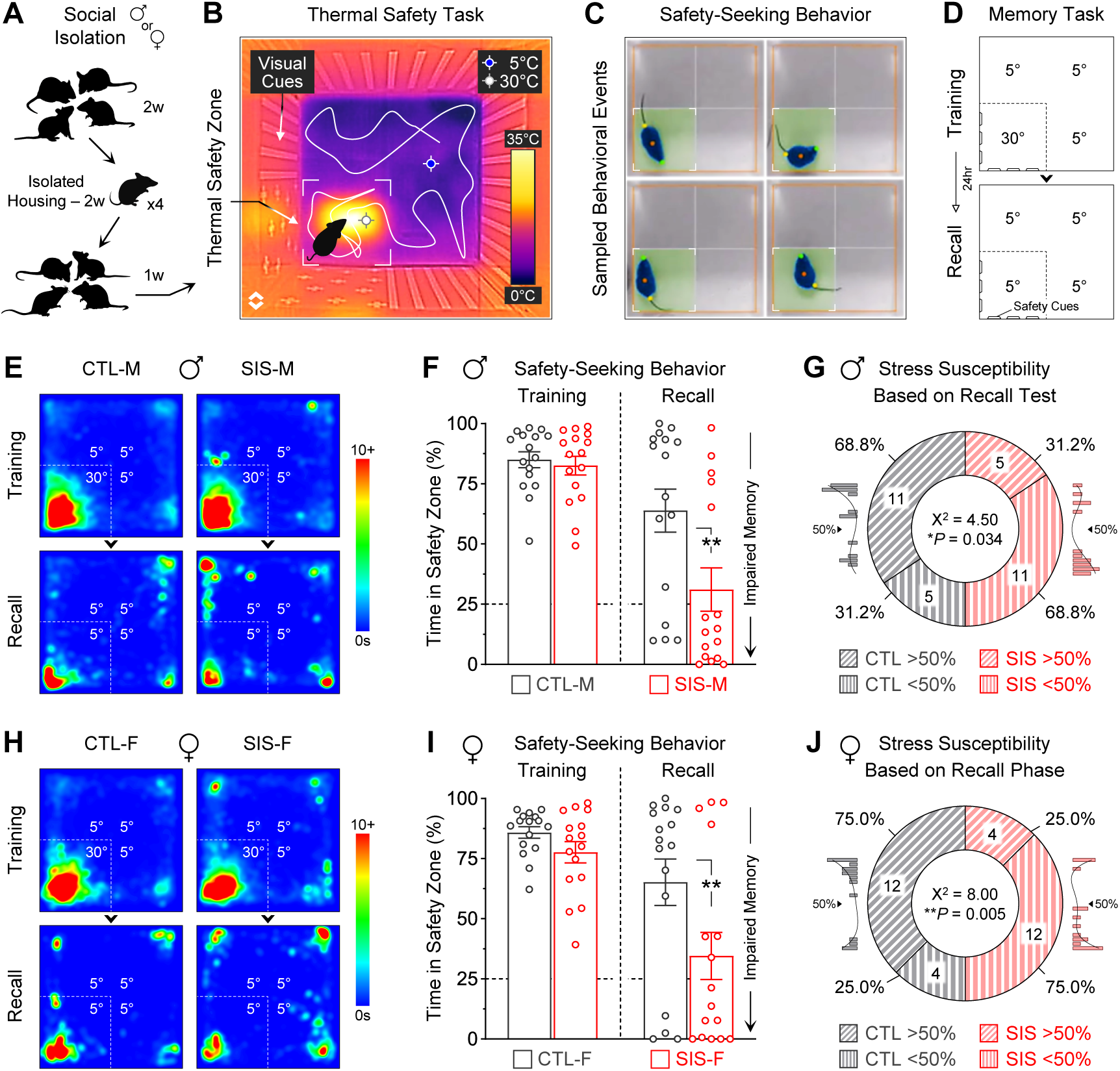
Social isolation stress led to robust impairment in safety memory, independent of mouse sex. **(A)** Depiction of a social isolation stress procedure. While controls remained in group-housing conditions, the experimental groups underwent single-housing for 14 days. After the isolation period, mice were returned to group-housing conditions with their former cagemates for a week, and then underwent training and testing in a safety learning task that involved a thermal threat. **(B)** Infrared image of the arena used for the thermal task, in which most quadrants had a noxious cold temperature (∼5°C, “threat zones”), while one quadrant had a pleasant warm temperature (∼30°C, “safety zone”). Visuospatial cues on the walls helped mice to differentiate the safety zone from the threat zones. (e.g., plus symbols vs vertical bars, counterbalanced across mice). **(C)** Examples of safety-seeking behavior. Software was used to track animals and quantify the time spent in the safety zone. **(D)** Experimental design to test for learning and memory. The training session lasted 10 min and included a warm safety zone, while the subsequent recall test lasted 5 min and no longer included the thermal reinforcer for the safety zone. **(E)** Average heatmaps for the male groups during the thermal task. **(F)** Quantifications of safety-seeking behavior for the male groups. Despite robust initial learning, many of the males that received the social isolation stress treatment exhibited significant memory impairment for the safety zone (\*\**P* = 0.0025). **(G)** Proportion of males that exhibited stress susceptibility or resiliency, based on memory recall. Consistent with the male distributions shown in the histogram insets, the cutoff was set to 50%. **(H)** Average heatmaps for the female groups during the thermal task. **(I)** Quantifications of safety-seeking behavior for the female groups. Similar to males, many of the females that received the social isolation stress treatment exhibited significant memory impairment for the safety zone (\*\**P* = 0.0086). **(J)** Proportion of females that exhibited stress susceptibility or resiliency, based on memory recall. Consistent with the female distributions shown in the histogram insets, the cutoff was set to 50%. [N = 16 per group; CTL, no-stress control; SIS, social isolation stress; M, males; F, females]

### Validation of the social isolation treatment as a significant stressor

The effectiveness of the two-week social isolation treatment as a stressor was evaluated in a subset of mice (N = 8 per sex per treatment, i.e., half of the controls and half of the stressed animals). Three methods of evaluation were implemented: (a) Changes in bodyweight gain, (b) Changes in exploration patterns during elevated-plus maze testing, and (c) Changes in locomotor activity during open-field testing. Bodyweights were assessed every other day during the social isolation period. The plus-maze and open-field assays were conducted twice, one day before and one day after the social isolation period. To minimize reductions in exploratory behavior due to decreases in contextual novelty, these assays were performed in counterbalanced manners in distinct procedure rooms.

The social isolation treatment significantly delayed the progression of bodyweight gain in both sexes. While the male groups exhibited a significant group × time interaction (Supp Figure 1B, Top Panel; *F*_(7,210)_ = 3.25, *P* = 0.0027), the female groups also exhibited a significant group × time interaction (Supp Figure 1B, Bottom Panel; *F*_(7,210)_ = 2.50, *P* = 0.018). In addition, the social isolation treatment produced significant reductions in open-arm exploration during plus-maze testing. While both male groups exhibited reductions when comparing the first and second plus-maze tests, only the reduction in the stress-male group was significant (Supp Figure 1C, Top Panel; Time Factor, *F*_(1,14)_ = 17.5, *P* = 0.0009; Post-Hoc Test for CTL-M, *P* = 0.21; Post-Hoc Test for SIS-M, *P* = 0.0054). Similarly, while both female groups exhibited reductions when comparing the first and second plus-maze tests, only the reduction in the stress-female group was significant (Supp Figure 1C, Bottom Panel; Time Factor, *F*_(1,14)_ = 15.2, *P* = 0.0016; Post-Hoc Test for CTL-F, *P* = 0.23; Post-Hoc Test for SIS-F, *P* = 0.011). Importantly, no significant alterations were observed in general locomotion during open-field testing for the male groups (Supp Figure 1D, Top Panel; Interaction Factor, *F*_(1,14)_ = 0.44, *P* = 0.52) or the female groups (Supp Figure 1D, Bottom Panel; Interaction Factor, *F*_(1,14)_ = 3.34, *P* = 0.89). Collectively, these results are consistent with well-established effects of stress, considering social isolation and other types of stressors, thereby validating once again the social isolation paradigm as a significant stressor that alters behaviors associated with internal states, such as anxiety (Walf and Frye, 2007; Mumtaz et al., 2018).

### Social isolation stress was detrimental for safety memory during the thermal task in both sexes

Males exhibited robust stress-induced impairments in safety memory during the thermal task. During the training session, similar to the control-male group, the stress-male group showed high levels of safety-seeking behavior (Figure 1E and Figure 1F; Training Session, CTL-M vs SIS-M, 85% vs 83% time spent in the safety zone; *P* = 0.80). However, during the subsequent recall test, while the control-male group showed relatively high levels of safety-seeking behavior, the stress-male group showed significantly lower levels of safety-seeking behavior (Figure 1E and Figure 1F; Recall Session, CTL-M vs SIS-M, 64% vs 31% time spent in the safety zone; *P* = 0.0025). Additional analysis of the individual data revealed that while many control-males showed good safety recall (11/16, 69%), some control-males showed poor safety recall (5/16, 31%). In contrast, while many stress-males showed poor safety recall (11/16, 69%), some stress-males showed good safety recall (5/16, 31%). A categorical chi-square test comparing these ratios revealed a significant difference between the control-male and stress-male groups (Figure 1G; X^2^ = 4.50, *P* = 0.034). These findings are consistent with our previous study.

Females also exhibited robust impairments in safety memory during the thermal task. As in the male groups, the control-female and stress-female groups showed high levels of safety-seeking behavior during the training session (Figure 1H and Figure 1I; Training Session, CTL-F vs SIS-F, 86% vs 78% time spent in the safety zone; *P* = 0.43). Despite this, during the subsequent recall test, while the control-female group showed relatively high levels of safety-seeking behavior, the stress-female group showed significantly lower levels of safety-seeking behavior (Figure 1H and Figure 1I; Recall Session, CTL-F vs SIS-F, 65% vs 35% time spent in the safety zone; *P* = 0.0086). Furthermore, the individual data revealed subsets of control-females showing good safety recall (12/16, 75%), subsets of control-females showing poor safety recall (4/16, 25%), subsets of stress-females showing poor safety recall (12/16, 75%), and subsets of stress-females showing good safety recall (4/16, 25%). A categorical chi-square test comparing these ratios revealed a significant difference between the control-female and stress-female groups (Figure 1J; X^2^ = 8.00, *P* = 0.005). Thus, the social isolation treatment produced similar susceptibility in males and females for the formation of lasting representations of safety.

### Active defensive mechanisms related to escape seemed unaffected by social isolation stress

Our next goal was to determine whether the stress treatment affected other behaviors that could potentially explain the impairments observed during the task. After inspecting a few representative videos, we realized that in addition to safety-seeking behavior, mice often exhibited multiple behaviors related to escape, including rearing, darting, and jumping. These behaviors represent active defensive mechanisms that help animals avoid harm as they experience threat (Bolles, 1969; Claudi et al., 2021). Therefore, a scoring method was implemented to quantify these behaviors for all animals and sessions and to perform comparisons across the control, stress, male, and female groups.

Rearing behavior was defined as periods in which mice adopted an upright standing posture, supporting their body on the hindlimbs while touching the sidewalls of the box with the forelimbs (Figure 2A). Rearing was the most frequent escape-related behavior during both phases of the thermal task (Figure 2B). While the rate of rearing behavior did not differ between the control and stress groups (stats in graph), males exhibited significantly more rearing behavior than females, but only during the recall test (MvsF; Training, *P* = 0.18; Recall, *P* = 0.015).

**Figure 2.**
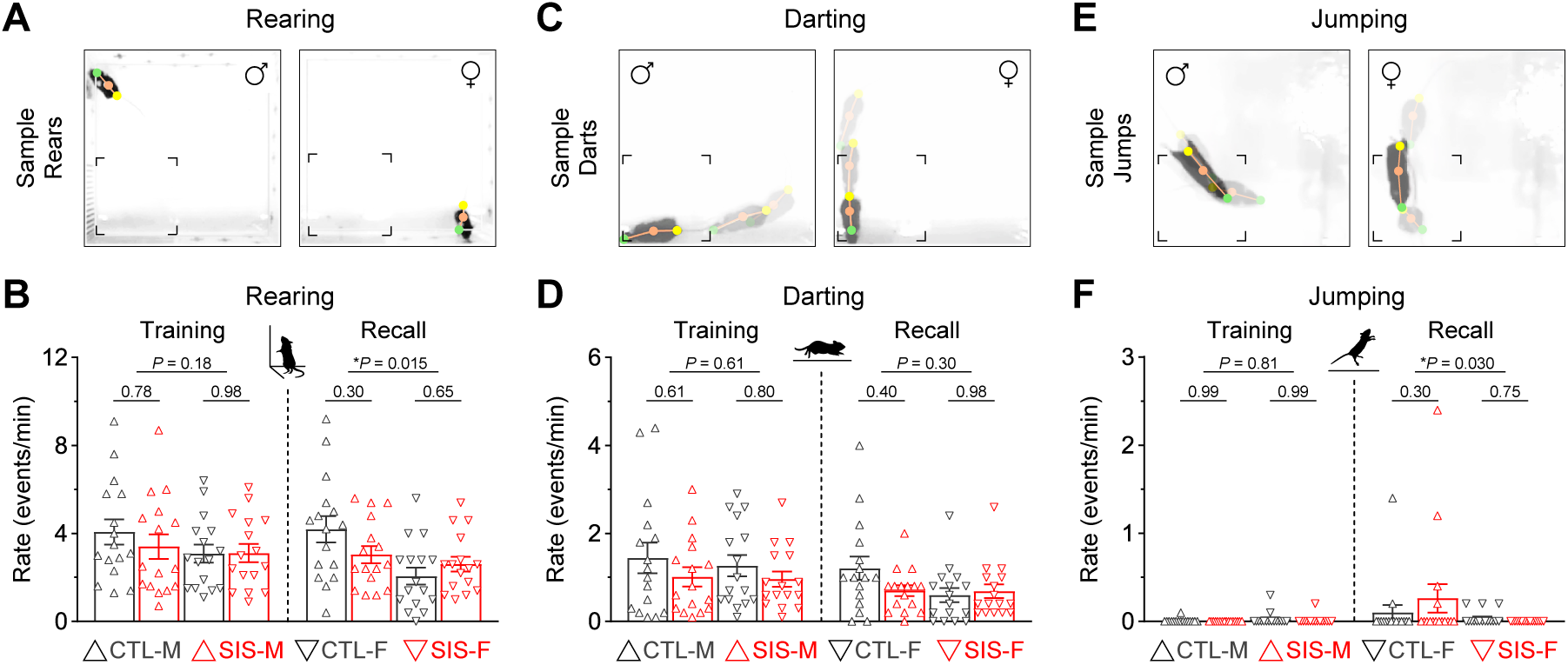
Active defensive mechanisms related to escape were strongly exhibited during the thermal safety task. **(A)** Rearing behavior, quantified when mice adopted an upright standing posture with forelimbs touching the sidewalls of the box. **(B)** Rearing behavior did not differ between the control and stress groups. However, males exhibited significantly more rearing than females during the recall test (\**P* < 0.05). **(C)** Darting behavior, defined as events in which mice made sudden flight-like running movements in any direction. Representative events are shown as overlaid video frames. **(D)** The rate of darting behavior did not differ between the control and stress groups, or the male and female cohorts. **(E)** Jumping behavior, defined as events in which mice exhibited sudden hopping or leaping movements. Representative events are shown as overlaid video frames. **(F)** While only a handful of mice exhibited jumping behavior, no significant differences were detected when comparing the control and stress groups. However, males tended to do more jumping than females, particularly during the recall test (\**P* < 0.05). [N = 16 per group; CTL, no-stress control; SIS, social isolation stress; M, males; F, females]

Darting behavior was defined as mice making sudden flight-like running moves in any direction from anywhere in the box (Figure 2C). While not as frequent as rearing, most animals exhibited some degree of darting, regardless of sex or treatment during both phases of the experiment (Figure 2D). The rate of darting behavior did not differ between the control and stress groups (stats in graph). Darting also did not differ between the male and female cohorts (MvsF; Training, *P* = 0.61; Recall, *P* = 0.30).

Jumping behavior was defined as mice performing sudden hopping or leaping moves anywhere in the box (Figure 2E). This was the least frequent escape-related behavior and was primarily exhibited by only a few animals (Figure 2F). While jumping was unaffected by stress (stats in graph), males tended to display more jumping behavior than females during the recall test (MvsF; Training, *P* = 0.81; Recall, *P* = 0.03). Thus, these active defensive mechanisms related to escape seemed to be unaffected by the stress treatment, despite some minor (*yet statistically significant*) sex differences.

### Passive defensive mechanisms also seemed unaffected by social isolation stress

Next, we evaluated possible stress-induced alterations in passive defensive mechanisms. This included behaviors such as freezing, crouching, and stretched postures. These behaviors have been associated with various functions, including protection, coping, and ambivalence during threat and adversity (Blanchard and Blanchard, 1969; Blanchard et al., 1986; Kaesermann, 1986).

Freezing behavior in mammals offers protection against threat in the environment (LeDoux and Daw, 2018). In this study, freezing behavior was defined as periods of minimal mobility for at least half a second (Figure 3A), which is consistent with other studies implementing quantifications for this behavior (Anagnostaras et al., 2010; Burgos-Robles et al., 2017). This behavior was the most frequent passive mechanism exhibited by animals during the thermal task (Figure 3B). While the rate of freezing episodes did not differ between the control and stress groups (stats in graph), males tended to exhibit more freezing behavior than females during both phases of the experiment (MvsF; Training, *P* = 0.031; Recall, *P* = 0.023). Sex differences in freezing behavior are consistent with previous observations during other behavioral tasks (e.g., Gruene et al., 2015; but see Borkar et al., 2020).

**Figure 3.**
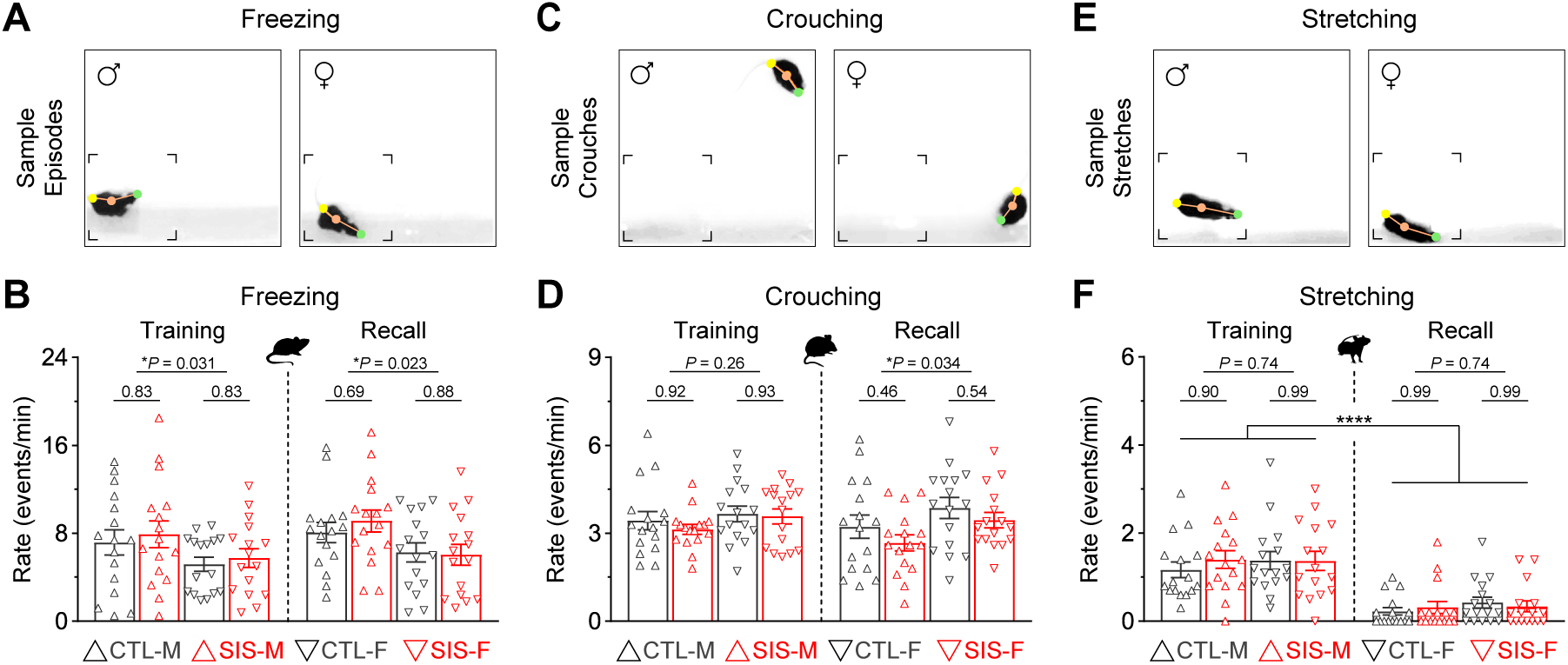
Passive defensive mechanisms related to protection, coping, and ambivalence were also prominent during the thermal safety task. **(A)** Freezing behavior, defined as periods of minimal mobility. This behavior is often implemented as a protection mechanism against threat, pain, or punishment. **(B)** The rate of freezing behavior did not differ between the control and stress groups. However, males exhibited significantly more freezing than females during both sessions (\**P* < 0.05). **(C)** Crouching behavior, defined as periods in which mice adopted a hunched posture with the head and forelimbs rolled inwards while supporting most of the body on the hindlimbs. This behavior is often implemented as a coping mechanism against cold temperatures. **(D)** The rate of crouching behavior did not differ between the control and stress groups. However, females exhibited significantly more crouching than males during the recall test (\**P* < 0.05). **(E)** Body stretching, defined as periods in which mice exhibited an elongated body posture. Stretched postures in rodents represent periods of ambivalence during threat, conflict, or uncertainty. **(F)** Body stretching was exhibited more prominently during the training session in all groups (\*\*\*\**P* < 0.0001). Yet, this behavior did not differ between the control and stress groups, or the male and female cohorts, during either training or recall. [N = 16 per group; CTL, no-stress control; SIS, social isolation stress; M, males; F, females]

Crouching behavior was defined as periods in which mice adopted a hunched posture with the head and forelimbs rolled inwards while supporting most of the body on the hindlimbs (Figure 3C). Warm-blooded animals often implement this behavior as a coping mechanism to regulate body temperature during exposure to cold (Chappell and Holsclaw, 1984). Mice of both sexes exhibited crouching behavior quite frequently during the thermal task (Figure 3D). While the rate of crouching did not differ between the control and stress groups (stats in graph), females tended to exhibit more crouching than males during the recall test (MvsF; Training, *P* = 0.26; Recall, *P* = 0.034).

Stretched body postures were also exhibited during the thermal task (Figure 3E). This passive defensive mechanism was not as frequent as freezing or crouching but was more prominent during the training session (Figure 3F; Training vs Recall, *P* < 0.0001 for all groups). Yet, the rate of stretched postures was unaffected by stress (stats in graph), and no significant differences were observed between males and females (MvsF; Training, *P* = 0.74; Recall, *P* = 0.74). Thus, similar to the active mechanisms examined above, these passive defensive mechanisms seemed unaltered by the stress treatment despite minor (yet statistically significant) differences between males and females.

### Other behaviors of interest also seemed insensitive to the social isolation stress treatment

Additional behaviors of interest during the thermal task were also evaluated for potential impact by stress. These included risk-taking, self-grooming, and general locomotion. Changes in risk-taking could represent alterations in decision-making processes (Laviola et al., 2003; Dent et al., 2014; Friedman et al., 2017). Self-grooming in rodents has been associated with many biological functions beyond cleanliness, including pain relief (Vos et al., 1998) and thermoregulation (Thiessen, 1988). Furthermore, alterations in self-grooming have been considered as behavioral markers of emotional distress, especially during situations involving conflict (Tinbergen, 1952; Rojas-Carvajal and Brenes, 2020). Changes in locomotion could represent alterations in mood, affect, or exploratory behavior (Yan et al., 2010; Seibenhener and Wooten, 2015).

Risk-taking during the thermal task was defined as periods in which mice protruded their head into the cold zones while keeping the rest of the body inside the warm zone (Figure 4A). This behavior was notably more prominent during the training session (Figure 4B; Training vs Recall, *P* < 0.0001 for all groups). While this behavior did not differ between control and stress (stats in graph), male mice exhibited significantly more risk-taking than the female mice (MvsF; Training, *P* = 0.009; Recall, *P* = 0.08). This is consistent with previous observations in which male rodents are more willing to perform behaviors that involve higher risk, whereas female rodents are more risk-averse and prefer to behave in a manner that involve lower risk (Orsini et al., 2016; Orsini and Setlow, 2017; Gomes et al., 2022; Shi et al., 2023).

**Figure 4.**
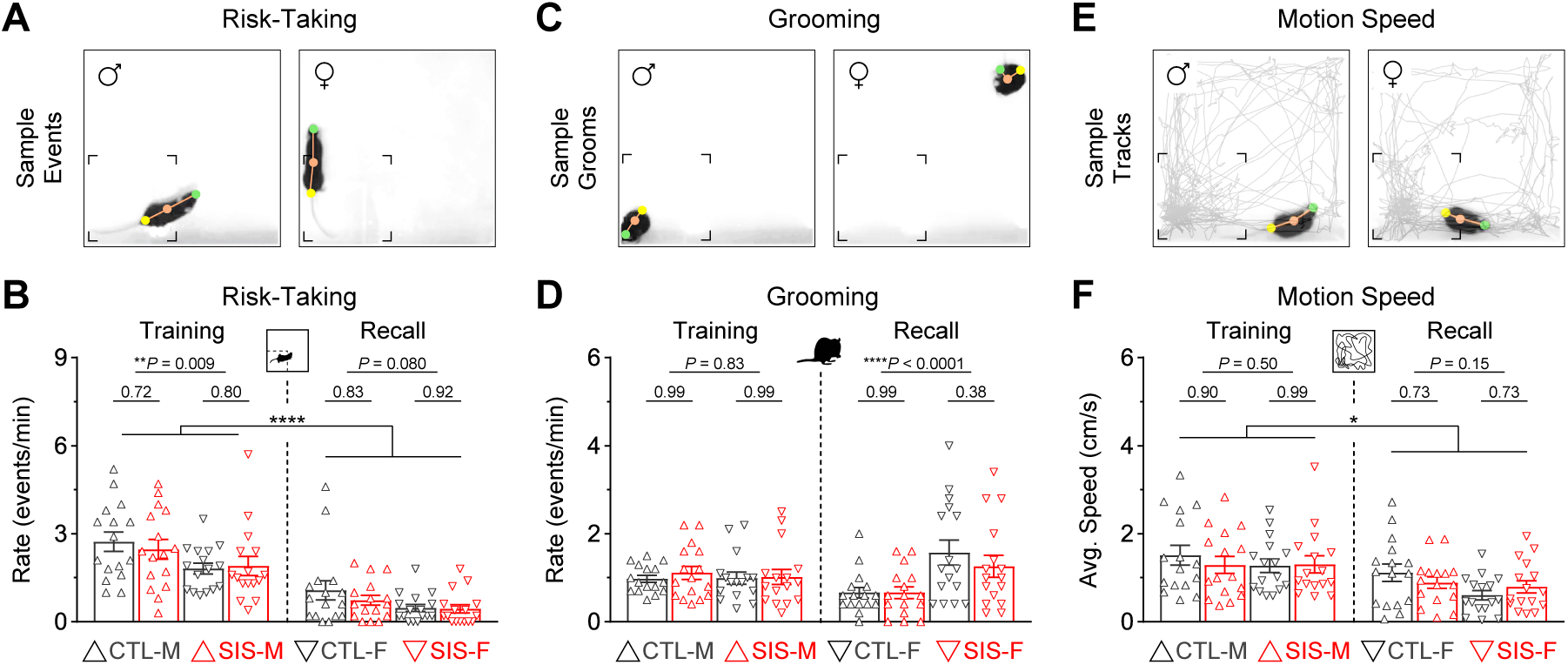
Other behaviors of interest during the thermal task. **(A)** Risk-taking behavior, defined as periods in which mice protruded their head into the cold zones while keeping the rest of the body inside the warm zone. **(B)** Risk-taking was exhibited more prominently during the training session in all groups (\*\*\*\**P* < 0.0001). While risk-taking did not differ between the control and stress groups, males exhibited significantly more risk-taking than females during the training session. **(C)** Grooming behavior as an index of stress during exposure to the cold. **(D)** Grooming rates did not differ between the control and stress groups. However, females exhibited a lot more grooming than males during the recall test (\*\*\*\**P* < 0.0001). **(E)** Motion speed as an index of general locomotion. **(F)** Although all the groups exhibited slightly faster average speeds during the training session than the recall test (\**P* < 0.05), the motion speeds did not differ between the control and stress groups, or the male and female cohorts. [N = 16 per group; CTL, no-stress control; SIS, social isolation stress; M, males; F, females]

Grooming was quantified when mice performed stereotyped behavior related to self-cleaning, including licking of the forelimbs, nose, face, head, or other parts of the body (Figure 4C). Grooming sequences were exhibited in similar amounts during both phases of the task (Figure 4D). While the rate of grooming events did not differ between the control and stress groups (stats in graph), females exhibited significantly more grooming behavior than males during the recall test (MvsF; Training, *P* = 0.83; Recall, *P* < 0.0001). This is inconsistent with prior studies showing that male rodents exhibit more self-grooming behavior than females during novelty or threat tasks (Thor et al., 1988; Borkar et al., 2020).

Finally, general locomotion was defined as the average speed of motion during each session (Figure 4E). All groups showed slightly higher motion speeds during the training session (Figure 4F; Training vs Recall, *P* < 0.05 in all groups). However, motion speed did not differ between the control and stress groups (stats in graph), or between the male and female groups (MvsF; Training, *P* = 0.50; Recall, *P* = 0.15). Therefore, similar to the other defensive mechanisms examined above, these additional behaviors seemed unaltered by the stress treatment and could not explain the profound alterations observed in the formation of lasting representations for the safety zone.

### The social isolation stress treatment seemed to disorganize the defensive mechanisms

Despite the stress treatment not impacting the overall quantity of defensive behavioral episodes, the final objective of this study was to investigate whether stress altered potential interactions and relationships among the behaviors. To achieve this, we implemented pairwise linear regression analysis considering all of the recorded behaviors, except for jumping which showed very low frequencies and would have skewed the analysis due to undersampling. After generating the linear regressions, the correlation coefficients were plotted into correlation matrix heatmaps using red-shifted colors to represent positive correlations and blue-shifted colors to represent negative correlations (Figure 5). Based on sixteen pair-wise samples, correlation coefficients greater than ±0.50 corresponded to significance levels of at least *P* < 0.05.

**Figure 5.**
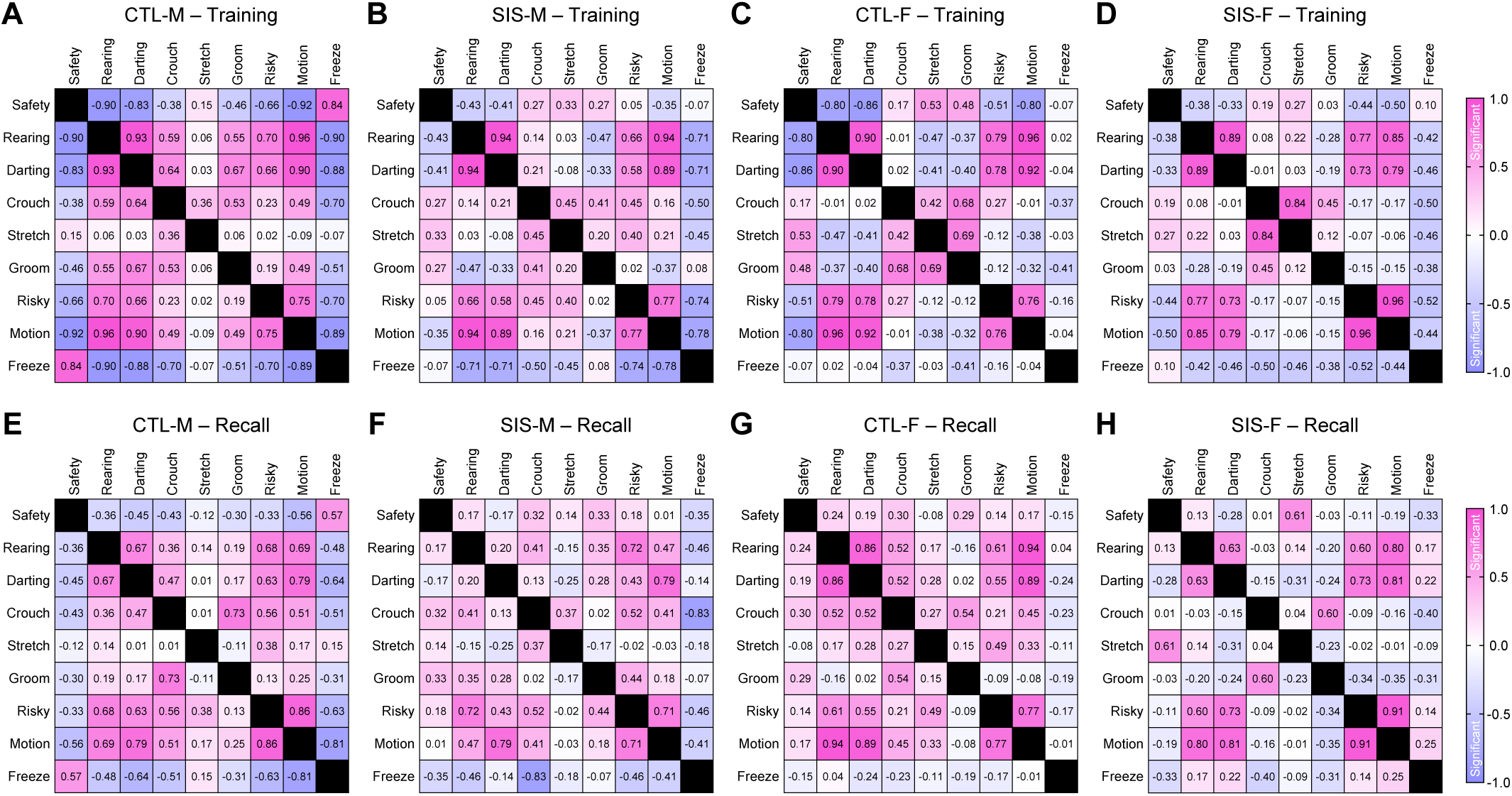
Linear regressions amongst the behaviors. **(A-D)** Regressions during the training session. **(E-H)** Regressions during the recall test. Correlations were computed for all the behavioral pairs, except for jumping behavior due to undersampling. These analyses considered gaussian distributions, sixteen sample per correlation, and two-tailed *P*-values. Correlation coefficients greater than ±0.50 corresponded to significance values of *P* < 0.05. Correlation coefficients greater than ±0.62 corresponded to significance values of *P* < 0.01. Correlation coefficients greater than ±0.75 corresponded to significance values of *P* < 0.001. Finally, correlations greater than ±0.82 corresponded to significance values of *P* < 0.0001. [CTL, no-stress control; SIS, social isolation stress; M, males; F, females]

In males, various significant relationships were detected between safety-seeking and other defensive mechanisms, especially during the training session in the control group but not in the stress group. For instance, safety-seeking was negatively correlated with rearing, darting, and risk-taking in the control-male group (Figure 5A; Safety vs Rearing, R = -0.90, *P* < 0.0001; Safety vs Darting, R = -0.83, *P* < 0.0001; Safety vs Risk-Taking, R = -0.66, *P* = 0.006). In contrast, these behaviors were uncorrelated in the stress-male group (Figure 5B; Safety vs Rearing, R = -0.43, *P* = 0.095; Safety vs Darting, R = -0.41, *P* = 0.12; Safety vs Risk-Taking, R = 0.05, *P* = 0.86). In addition, safety-seeking was positively correlated with freezing behavior in the control-male group (Figure 5A; Safety vs Freezing, R = 0.84, *P* < 0.0001), whereas these behaviors were uncorrelated in the stress-male group (Figure 5B; Safety vs Freezing, R = -0.07, *P* = 0.81). Thus, the stress treatment seemed to affect the organization of these behaviors.

Similar to males, the female groups exhibited significant correlations of safety-seeking, rearing, darting, and risk-taking during training in the control group but not in the stress group. That is, safety-seeking was negatively correlated with rearing, darting, and risk-taking in the control-female group (Figure 5C; Safety vs Rearing, R = -0.80, *P* = 0.0002; Safety vs Darting, R = -0.86, *P* < 0.0001; Safety vs Risk-Taking, R = -0.51, *P* = 0.044). In contrast, these behaviors were uncorrelated in the stress-female group (Figure 5D; Safety vs Rearing, R = -0.38, *P* = 0.15; Safety vs Darting, R = -0.33, *P* = 0.21; Safety vs Risk-Taking, R = -0.44, *P* = 0.09). In contrast to males, safety-seeking and freezing behavior were not correlated in either the control-female group (Figure 5C; Safety vs Freezing, R = -0.07, *P* = 0.79) or the stress-female group (Figure 5D; Safety vs Freezing, R = 0.10, *P* = 0.73). Such sex difference could be attributed to the observation of significantly lower freezing rates in females than males.

Unlike the training session, during the memory recall test, safety-seeking showed limited correlations with the other defensive behaviors in all groups (Figures 5E-H). Moreover, some behaviors exhibited very stable relationships across all groups during both sessions (e.g., rearing and darting). Finally, crouching and stretched postures showed very limited correlations or did not exhibit clear patterns of effects by stress during either of the sessions. Collectively, these findings suggest that stress may have reorganized specific defensive behaviors, such as rearing, darting, and risk-taking, in a manner that mice were still capable of exhibiting these behaviors, but they were exhibited in chaotic manners in relationship to safety-seeking. Disorganized defensive behavior during training could represent a significant mechanism by which stress led to alterations in the formation of safety memory.

## Discussion

This study evaluated the negative impact of social isolation stress, as well as potential sex differences in mice during a behavioral task that involved memory formation for a spatial zone that predicted thermal safety. In addition to the impact of stress on memory formation, major focus was placed on to the assessment of stress-induced alterations in a variety of behaviors, including active defensive mechanisms (e.g., rearing, darting, and jumping), passive defensive mechanisms (e.g., freezing, crouching, and ambivalent posture), and other relevant behaviors (e.g., risk-taking, grooming, and general locomotion). This represents a broad and novel approach from which the results obtained in this study provide new insights into the mechanisms that potentially contribute to the impacts of stress on safety learning. Overall, our results showed that lasting representations of safety are highly susceptible to social isolation stress in both male and female mice. Furthermore, while the stress treatment did not affect the average frequency in which animals of either sex exhibited the defensive behaviors listed above, the stress treatment reorganized the structure of some of those behaviors so that they became completely uncorrelated with safety-seeking behavior. These findings highlight the severity of social isolation as a critical stressor that alters behavioral mechanisms that are critical for adaptability, resiliency, and safety learning when animals experience threat and adversity.

### Advantages for evaluating safety learning with a spatial-based task

Safety learning has been traditionally evaluated using “conditioned inhibition paradigms” in which a particular cue that explicitly omits a transient noxious stimulus (typically an electric footshock) is capable of diminishing fear-related responses to another cue that explicitly predicts the delivery of the noxious stimulus (Kendrick, 1958; Odriozola and Gee, 2021). While improvements have been made over the years, some issues remain unresolved for the conditioned inhibition paradigms. For instance, while threat memory is clearly represented by a defensive behavioral response such as freezing, safety memory is not defined by a particular behavior, but safety is instead inferred from the reductions observed in freezing behavior when the threat and safety cues are presented simultaneously (Donahoe and Palmer, 1988; Sosa and Ramírez, 2019). In contrast, this is not an issue for the behavioral task used in the present study, which our group recently developed and validated (Felix-Ortiz et al., 2024). In our task, mice are able to learn safety-seeking behavior by exploring an open-field arena with thermal threat and thermal safety zones (5°C vs 30°C, respectively). Importantly, mice are allowed to use visual cues for spatial orientation, navigation, and learning about the particular zone that is associated with thermal safety. Recognition of the visual cues guide safety-seeking behavior during subsequent memory tests. These features render this task particularly suitable for evaluating the mechanisms of safety learning and long-term memory recall through the assessment of a clearly quantifiable behavior.

The present study further expands the idea of clearly quantifiable behavior during the thermal safety task by showing that a repertoire of defensive behaviors related to escape, protection, and risk-taking are also manifested in prominent manners during this paradigm. However, many of the observed defensive mechanisms seem to be related to innate reactions to the noxious cold temperature rather than learned behavioral reactions (Blanchard et al., 1986; LeDoux and Daw, 2018). Nonetheless, some behaviors, including darting, rearing, and risk-taking, correlated quite strongly with safety-seeking behavior in both sexes during the training session. Furthermore, the social isolation stress treatment rendered the relationships among those behaviors insignificant. This could represent an important mechanism by which the stress treatment rendered animals more susceptible to forgetting the safety memory. Yet, further investigation is needed to determine how each defensive behavior contributes to safety learning. For instance, neural pathways critical for particular behaviors (e.g., from the dorsal peduncular prefrontal cortex to the central amygdala, which is required to produce flight-related escape responses; Borkar et al., 2024) could be individually targeted as mice undergo our thermal safety task. Approaches like this could reveal important roles for individual behaviors during safety learning.

### Sex differences in threat-related behavior and safety learning

During the thermal task, males exhibited more freezing behavior than females during both the training and recall sessions. Furthermore, males also exhibited more rearing, jumping, and risk-taking behavior than females, whereas females exhibited more crouching and grooming than males during some sessions. These findings are consistent with growing evidence that male and female rodents could implement distinct behavioral strategies when dealing with situations involving threat. For instance, during classical threat conditioning (i.e., cue predicts shock), while freezing behavior characterizes fear-related responses in males, females preferably exhibit darting behavior (Gruene et al., 2015; Clark et al., 2019; Mitchell et al., 2022). In contrast, newer versions of threat conditioning that allow transitions from freezing to darting have yielded observations that are somewhat contradictive, as males showed less freezing behavior than females (Borkar et al., 2020). Despite this, differences in the learning contingencies during distinct tasks could explain the discrepancies in behavioral sex differences. Nonetheless, future studies could implement approaches in which male and female animals undergo systematic testing in a variety of behavioral tasks to better understand sex-specific behavioral differences.

There is also some documented evidence for sex differences in safety learning, which has been mostly gathered using cue discrimination and/or the conditioned inhibition strategy. For example, one study showed that while female rats discriminated between a threat cue and a safety cue better than males, females did not exhibit better suppression of the fear response than males when the cues were presented simultaneously (Foilb et al., 2018). Furthermore, another study showed that while male rats were capable of reducing fear responses when the threat and safety cues were presented simultaneously, female failed to exhibit significant reductions in fear responding (Krueger and Sangha, 2021). In addition, another study showed that similar to male rats, females in metestrus/diestrus (i.e., low-estrogen phase) were capable of discriminating well between the threat and safety cues, whereas females in proestrus (i.e., high-estrogen phase) exhibited poor discrimination among the cues (Trask et al., 2020). Thus, while contradictive, some of these discrepancies could have resulted from differences in phases of the estrous cycle, as well as potential differences in threat-related behaviors other than freezing. Notably, this field of behavioral neuroscience has evolved enough to realize that threat and safety learning are not only commanded by freezing responses but also that male and female animals are capable of implementing distinct strategies when coping with situations involving threat (Shansky, 2024). While the present study did not consider the estrous cycle, future studies could implement such measurements to evaluate the possible contribution of sex-specific hormonal milieus during the thermal safety task (Peyrot et al., 2020).

### Sex differences in the influence of stress on threat-related behavior

Growing evidence suggests that females are more susceptible to the impact of stress than males. For instance, female animals are more prone to bodyweight loss and the development of hyperactive, anxious-like behavior with chronic stress (Furman et al., 2022). Furthermore, some studies have reported that stress-exposed females are more susceptible to acquiring stronger threat memories that are more resistant to extinction than sex-matched, no-stress controls (Baran et al., 2009; Sanders et al., 2010). While these observations are consistent with the greater risk for stress and trauma-related disorders observed in human females (Kessler et al., 2012; Gradus, 2017), the evidence presented above, as well as in many other studies (reviewed in Farrell et al., 2013; Peyrot et al., 2020) primarily considers the traditional views of stress-induced alterations in the acquisition and/or extinction of threat memory. However, much less is known about the impact of stress in safety learning or whether safety learning in female animals could be more impacted by stress than in male animals.

### Concluding remarks

Despite the fact that stress produced similar effects in males and females, this study represents a significant step forward toward developing a better understanding of the impacts of stress and sex factors in safety learning. After considering at least ten behaviors during the safety task, our female subjects did not exhibit a greater impact of stress than the male subjects. Furthermore, additional measurements during the plus-maze and open-field tasks also revealed no greater impacts of stress in the female subjects. Nonetheless, we only evaluated social isolation stress, and future studies could implement other types of stressors to either refute our results or confirm that female mice are indeed similarly vulnerable to stress than male mice with respect to safety learning.

## Acknowledgements

The authors thank other members of the Burgos Lab for feedback on this project. The project was supported by the College of Science at the University of Texas at San Antonio, University of Texas Program for Science and Technology Acquisition and Retention, and Baptist Health Foundation.

## Author Contributions

S.A.V., J.R.M., and A.B.R. designed the experiments. S.A.V., A.C.F.O., and K.L.O. performed the experiments. S.A.V. and A.B.R. analyzed the data and prepared the manuscript.

## Supplementary Figure Legend

**Supp Figure 1.**
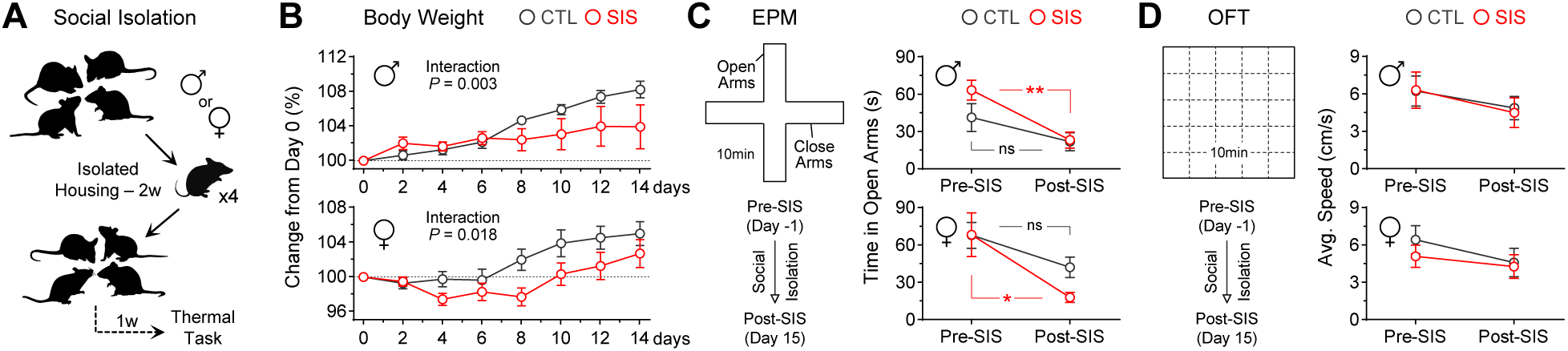
Validation of the social isolation procedure. **(A)** Depiction of timeline of the social isolation stress procedure. **(B)** Bodyweight measurements every other day during the social isolation procedure. Delays in bodyweight gain represent effectiveness of treatments as emotional stressors. **(C)** Behavioral testing in the elevated-plus maze (EPM) before and after the social isolation procedure. Significant decays in the time of open arm exploration represent effectiveness of treatments as emotional stressor. While the no-stress control groups showed no significant changes between the pre and post sessions (Males, *P* = 0.21; Females, *P* = 0.23), the stress groups exhibited significant decays in open arm exploration (Males, \*\**P* = 0.0054; Females, \**P* = 0.011). **(D)** Traditional open-field test (OFT) before and after the social isolation procedure. No significant changes were detected for the average speed of motion in the open field (all *P’s* > 0.53). [N = 8 per group CTL, control; SIS, stress]

